# The combinatorics of discrete time-trees: theory and open problems

**DOI:** 10.1101/063362

**Authors:** Alex Gavryushkin, Chris Whidden, Frederick A Matsen

**Author notes:** CW and FM were funded by National Science Foundation awards 1223057 and 1564137. CW is a Simons Foundation Fellow of the Life Sciences Research Foundation. FM supported in part by a Faculty Scholar grant from the Howard Hughes Medical Institute and the Simons Foundation.

## Abstract

A *time-tree* is a rooted phylogenetic tree such that all internal nodes are equipped with absolute divergence dates and all leaf nodes are equipped with sampling dates. Such time-trees have become a central object of study in phylogenetics but little is known about the parameter space of such objects. Here we introduce and study a hierarchy of discrete approximations of the space of time-trees from the graph-theoretic and algorithmic point of view. One of the basic and widely used phylogenetic graphs, the NNI graph, is the roughest approximation and bottom level of our hierarchy. More refined approximations discretize the relative timing of evolutionary divergence and sampling dates. We study basic graph-theoretic questions for these graphs, including the size of neighborhoods, diameter upper and lower bounds, and the problem of finding shortest paths. We settle many of these questions by extending the concept of graph grammars introduced by Sleator, Tarjan, and Thurston to our graphs. Although time values greatly increase the number of possible trees, we show that 1-neighborhood sizes remain linear, allowing for efficient local exploration and construction of these graphs. We also obtain upper bounds on the *r*-neighborhood sizes of these graphs, including a smaller bound than was previously known for NNI.

Our results open up a number of possible directions for theoretical investigation of graph-theoretic and algorithmic properties of the time-tree graphs. We discuss the directions that are most valuable for phylogenetic applications and give a list of prominent open problems for those applications. In particular, we conjecture that the split theorem applies to shortest paths in time-tree graphs, a property not shared in the general NNI case.

## 1. Introduction

The last ten years have seen an explosion of methods using sequence data to infer demographic model parameters by sampling phylogenetic trees (Kuhner, Yamato, and Felsenstein 1995; Kuhner, Yamato, and Felsenstein 1998; Kuhner, Beerli, et al. 2000; Beerli and Felsenstein 2001; Kuhner 2006; Drummond, Nicholls, et al. 2002; Drummond, Rambaut, et al. 2005; Drummond, Ho, et al. 2006; Minin, Bloomquist, and Suchard 2008). These methods have had an especially significant impact in the study of quickly evolving organisms such as viruses, such as inferring historical epidemic spreading rates. For example, such methods can be used to infer time to a most recent common ancestor of human HIV group M viruses (Worobey et al. 2008; Baele et al. 2013). Thus the *time-tree*—a rooted phylogenetic tree with all internal nodes equipped with absolute divergence dates— has become an important object of investigation. This interpretation of continuous parameters for time-trees stands in contrast to that for classical phylogenetic trees, in which branch lengths quantify the amount of molecular substitution along a branch. Posterior distributions on both time-trees and classical trees are estimated using Markov chain Monte Carlo (MCMC) (Mau and Newton 1997; Yang and Rannala 1997; Drummond, Nicholls, et al. 2002), but with different transition kernels. This paper builds a foundation for a mathematical understanding of time-trees, such as understanding convergence properties of MCMC thereupon.

Although Markov chain Monte Carlo (MCMC) is guaranteed to sample from the true posterior given an infinite run time, it is important to understand mixing properties of the chain, which determine sampling properties for a finite time run. The mixing properties of phylogenetic MCMC have, in the classical unrooted case, been a major area of research from both the theoretical perspective of mixing time bounds (Mossel and Vigoda 2005; Mossel and Vigoda 2006; Štefankovič and Vigoda 2011; Spade, Herbei, and Kubatko 2014) and the practical perspective of performance on real data (Beiko et al. 2006; Ronquist et al. 2006; Lakner et al. 2008; Whidden and Matsen IV 2015). This research has focused on mixing over the set of discrete phylogenetic tree graph structures, because mixing over these discrete structures is the primary obstruction to MCMC convergence.

Time-trees use different MCMC transition kernels than do classical phylogenetic trees. The discrete component of MCMC moves between classical phylogenetic trees are typically determined by their discretization. For example, common moves include subtree *prune and regraft* (SPR) moves, which cut a subtree off and reattach it at another location, and the subset of SPR moves called nearest-neighbor *interchange* (NNI) moves. Classically, these are applied without reference to branch lengths. On the other hand, the discrete component of moves between time-trees are defined also in terms of their timing information. For example, Hohna, Defoin-Platel, and Drummond (2008) show that SPR-like moves that reattach subtrees at the same divergence time are more effective than ones that do not. This sort of move cannot be expressed using the type of discretization used thus far in which all continuous information is lost, and so the previous work on MCMC mixing cannot be applied.

However, one can discretize time-trees in a way that does preserve some of the information. For example, by retaining the order of internal nodes backward in time one obtains a so-called *ranked tree* (Semple and Steel 2003). To make a less rough approximation of a time-tree, one can allow the time periods between nodes of the tree to take only finitely many possible values (Åkerborg, Sennblad, and Lagergren 2008). This results in an object we call a *discrete time-tree*. The graph built on such trees provides a discretization of the space of time-trees, which we call a *discrete time-tree graph*. By a *graph* on a set of trees here and throughout the paper we mean a graph consisting of trees as vertices, with edges connecting pairs of trees that are identical after a given tree rearrangement operation (Semple and Steel 2003). This graph-theoretic terminology is convenient and has become widespread in recent years (Spade, Herbei, and Kubatko 2014; Whidden and Matsen IV 2015; Gavryushkin and Drummond 2016). A sequence of time-trees sampled using MCMC projects to a collection of movements on a graph in which each vertex is a discrete time-tree and each edge is a discrete version of an MCMC move on time-trees.

Although inferential algorithms have applied MCMC on time-trees for over a decade, and graphs corresponding to discretizations of unrooted tree space have been studied for even longer, we are not aware of any work defining graphs from discretization of time-tree spaces or analyzing random walks thereupon. Such a theory would provide a foundation for understanding the behavior of MCMC algorithms on time-trees, as has been done previously for graphs associated with unrooted phylogenetic trees. Up to now, the only discretization of time-tree space is that of ranked trees (Page 1991; Ford, Matsen, and Stadler 2009; Lambert and Stadler 2013), and the corresponding graphs have not been studied.

In this paper we initiate the mathematical study of discrete time-tree graphs, obtain basic geometric and graph-theoretic results, and compare these results to those in the classical phylogenetic setting. In particular, we focus on ranked trees, discrete time-trees (as defined above), and the ultrametric versions of those types of trees. We establish size bounds for neighborhoods and diameters in these tree spaces and show how those bounds can be used to develop efficient tree search algorithms such as MCMC. The importance of these basic geometric characteristics in phylogenetics and other areas of evolutionary biology is highlighted in (Huber et al. 2011).

## 2. Technical introduction

Throughout the paper by a (phylogenetic) *tree* we mean a rooted binary tree with designated leaves, that is, an undirected acyclic graph with the following properties: (1) all nodes have degree 1, 2, or 3; (2) there exists exactly one node of degree 2, this node is called the *root* of the tree; (3) all nodes of degree 1 are labeled by distinct identifiers, these nodes are called *leaves* or *taxa* (singular *taxon*). The *parent* of a node *x* in a tree is the unique node *y* that is both adjacent to *x* and closer to the root of the tree than *x*. Every node of a tree has a parent except for the root, and is called the *child* of that parent.

We fix the number of leaves of the trees and denote this number by *n* throughout the paper. We also assume that each tree with *n* leaves uses the same fixed set of *n* labels to mark the leaves. Hence, we say that two trees are *isomorphic* if they are isomorphic as graphs and the isomorphism maps leaves marked by the same label to each other. We do not distinguish between isomorphic trees, i.e. we *identify* them.

By a *time-tree* (Figure 1) we mean a phylogenetic tree with an absolute time associated with every node of the tree so that the time strictly increases along every path from a leaf to the root: for internal nodes the time is interpreted as (typically estimated) divergence time and for leaves it is the (typically known) sampling time. By “absolute” we clarify that these are not relative times, but rather actual times that can be put on a calendar. We assume that time progresses backward, from the leaves to the root.

**Figure 1.**
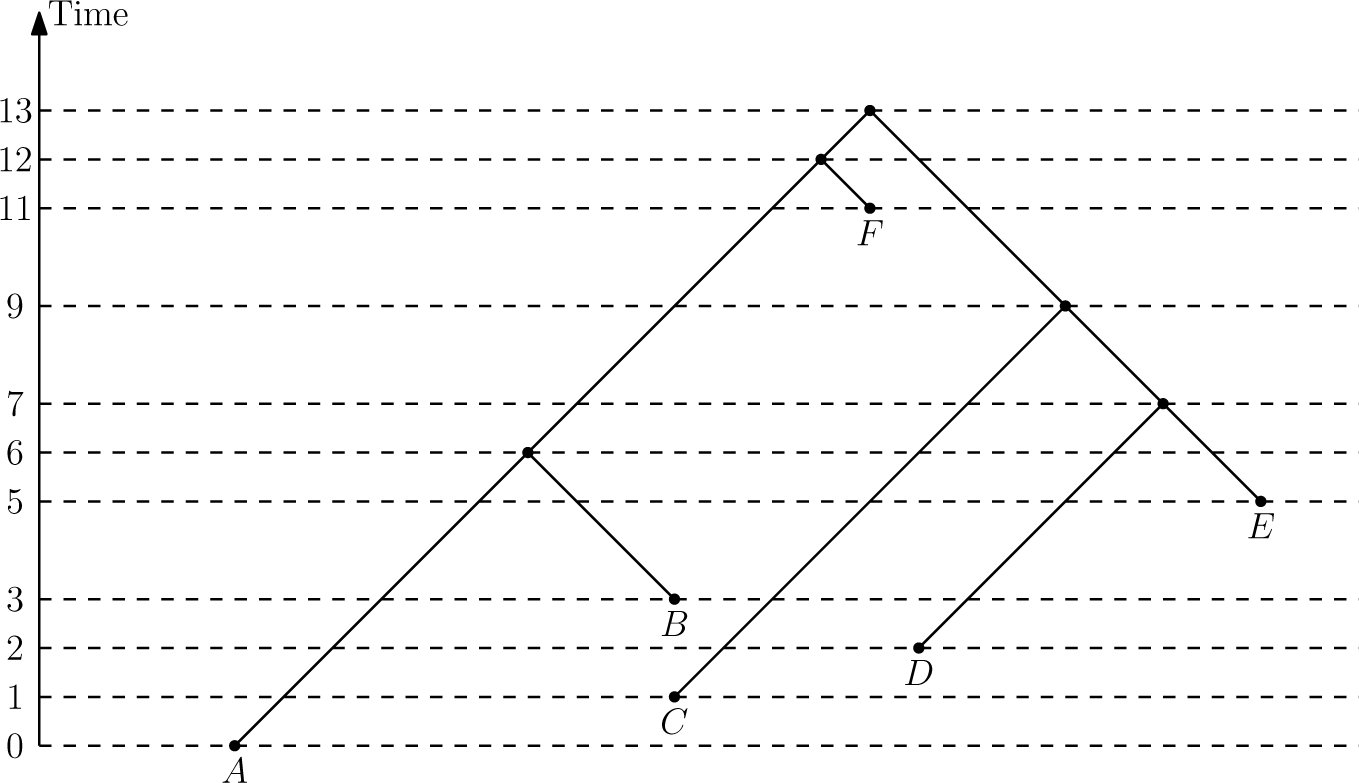
Time-tree on 6 leaves. Time is measured by non-negative real numbers. If all times are integers, the tree is a discrete time-tree.

A *discrete time-tree* is a tree such that all its nodes are assigned distinct times from the set of non-negative integers and every node has a smaller time than its parent (Figure 1). Note that this implies that for every pair of nodes *x, y* if the shortest path from the root to *x* passes through *y* then the time of *y* is greater than the time of *x*. Each node is associated with a type of event: internal nodes represent divergence events, while leaf nodes represent the sampling of new taxa. The *rank* of a node is the number of nodes in the tree with strictly smaller time. We say that a pair of nodes *x, y* of a discrete time-tree is an *event interval* if there exists no node *z* such that the time of *z* is between the time of *x* and *y*. Hence, an event interval is an interval between two divergence events, a taxon and a divergence event, or two taxa. Note that taxon events can be younger than divergence nodes, i.e. when the sampling time precedes divergence events on the tree. The difference between the times of *x* and *y* is called the *length* of the interval *x, y*. We identify two discrete time-trees if they are isomorphic as trees and the isomorphism preserves ranks of the nodes as well as event interval lengths.

We are now ready to introduce a hierarchy of discrete time-trees. At the bottom level of our hierarchy is the well-known NNI graph, which does not have timing information. NNI is a graph with the vertices being all trees on *n* leaves. Two trees *T* and *R* are adjacent in NNI if there exists an edge *e* in *T* and an edge *f* in *R* such that both edges are not adjacent to a leaf and the graph obtained from *T* by shrinking *e* to a vertex is isomorphic to the graph obtained from *R* by shrinking *f*. We denote this graph by DtT0, where DtT stands for “discrete time-trees”.

The following graph DtT_*m*_ forms the level *m* > 0 of the hierarchy. The set of vertices of the graph is the set of all discrete time-trees on *n* leaves such that every event interval has its length not greater than *m*. Two trees *T* and *R* are adjacent in DtT_*m*_ if *R* can be obtained from *T* by one of three operations: a *length move performed on interval I*, *swapping the rank of two nodes x and y on interval I* or an NNI *move performed on interval I* (Figure 2). A length move changes the length of *I* by 1. Swapping the rank of two nodes swaps the times of the nodes; such a swap is only possible when the nodes bound an event interval *I* of length 1. *R* can be obtained from *T* by an NNI move if there exist event intervals *I_T_* and *I_R_* of length 1 in *T* and *R*, respectively, such that the graphs obtained by shrinking *I_T_* and *I_R_* to vertices are isomorphic and the isomorphism preserves the lengths of the event intervals. In other words, by going from one tree to an adjacent tree in the DtT*_m_* graph we can either change the length of one event interval by one unit, swap the rank of two nodes bounding an event interval of minimal length, or send the length of an event interval of minimal length down to zero and then resolve the multifurcation to either of the two possible trees. In the latter case, the new interval is of minimal length. See Figure 3 for an example of the full variety of possible moves.

**Figure 2.**
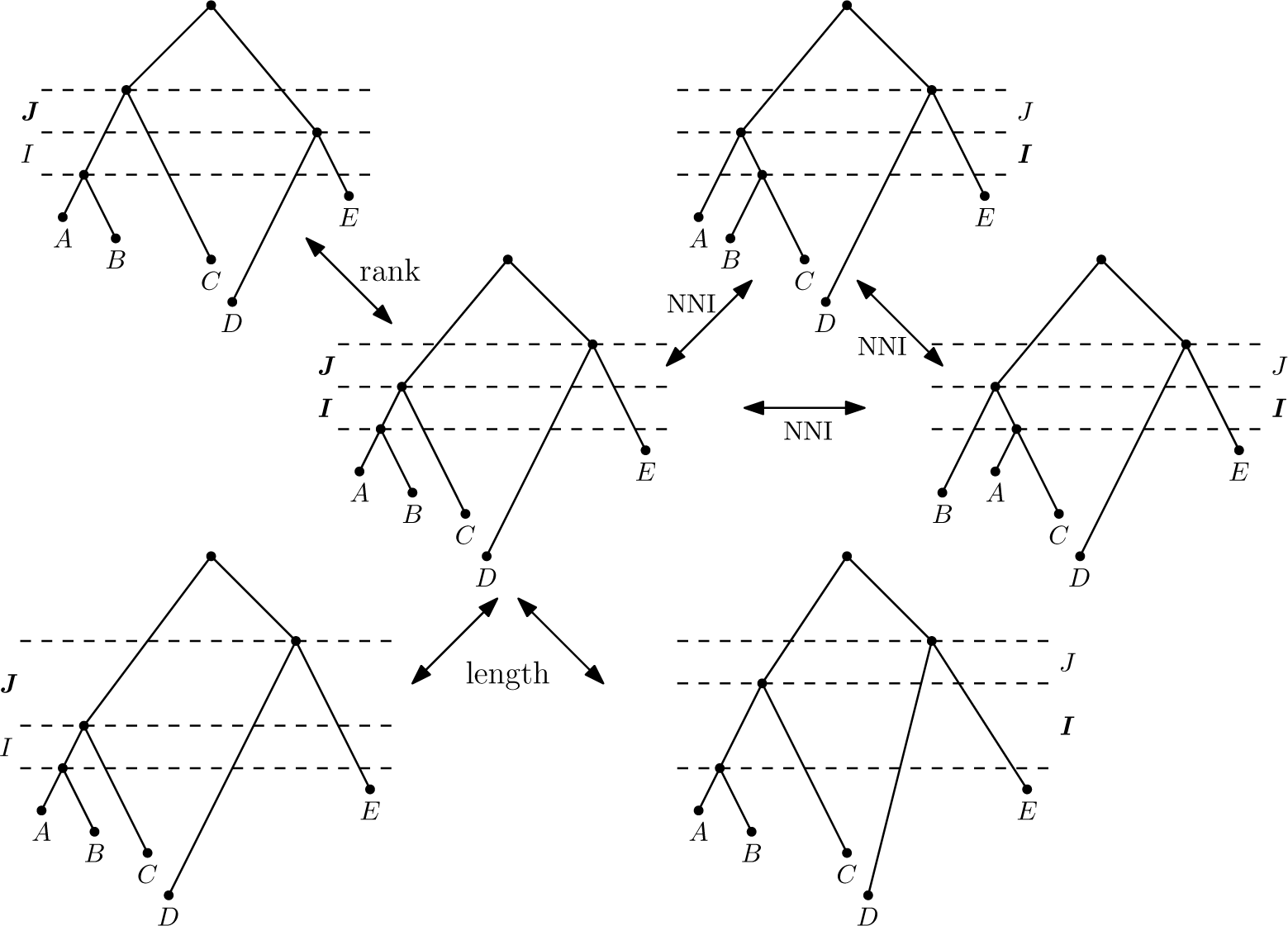
All possible moves performed on intervals *I* and *J*. Assuming that the length of every event interval is either 1 or 2, the outer trees are all possible neighbors in DtT of the tree in the middle obtained by moves performed on intervals *I* and *J*. The trees on the right are obtained by moves performed on interval *I* and those on the left on *J*.

**Figure 3.**
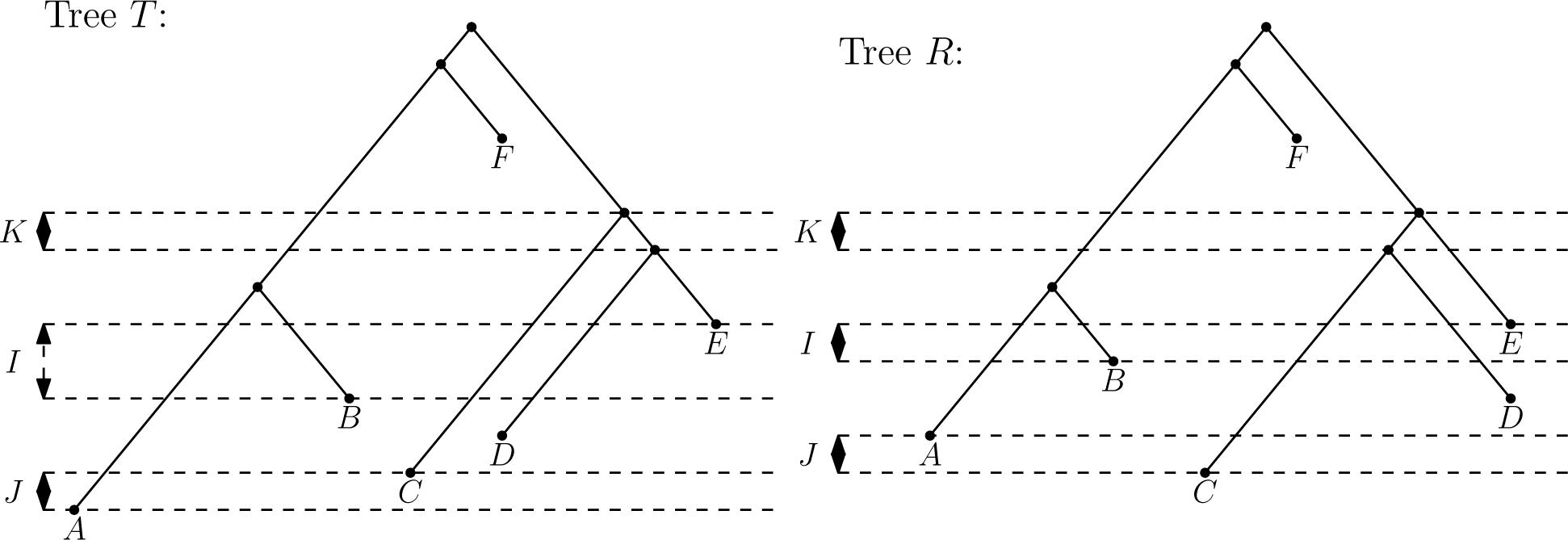
Trees *T* and *R* are at DtT_2_ distance 3. To move from *T* to *R* in DtT_2_, one must decrease the length of interval *I*, swap the ranks of nodes *A* and *C* bounding interval *J*, and perform an NNI move on interval *K*, resolving nodes *C* and *D*.

Note that DtT_0_ can be obtained from DtT_1_ by “forgetting” ranks and DtT_1_—from DtT*_m_* by forgetting lengths. In general, the graph structure for *m* > 1 can be understood by considering all four-tree diagrams as the four trees on the right in Figure 2 and introducing an extra layer of nodes in every tail adjacent to the NNI triangle.

This hierarchy can be seen as a set of discrete refinements of the full space of phylogenetic time-trees, where event intervals can take an arbitrary real value. Indeed, if we allow the lengths of event intervals to take every possible non-negative real value and impose the Euclidean metric on trees with the same ranked topology, then we get the *τ*-space introduced by Gavryushkin and Drummond (2016). In this case, DtT_1_ is the graph with the vertices being the orthants of *τ*-space and the adjacency relation being the relation of “have a shared facet of co-dimension 1.” Similarly DtT_0_ is the adjacency graph on the orthants of BHV space (Billera, Holmes, and Vogtmann 2001).

The DtT*_m_* graph on ultrametric trees, that is the set of trees such that each leaf has the same time, is denoted by DtTu*_m_*. The DtT1 graph is denoted by RNNI and called the *ranked* NNI graph; on ultrametric trees we will denote it RNNIu. The R in RNNI stands for *ranked* and not for rooted. We emphasize that although this is the RNNI *graph*, NNI *moves* can be applied to trees in any of the spaces considered (under appropriate conditions). For simplicity, we use the generic identifier DtT to refer to a DtT*_m_* with an arbitrary *m* > 1.

A graph can also be seen as a metric space where the distance is given by the length of a shortest path, so we will refer to DtT_m_ and other graphs introduced below as both, in agreement with (Semple and Steel 2003).

## 3. Geometry and complexity of discrete time-trees

We begin by considering the shortest path distance on the graphs. It is well-known that computing distances is NP-hard in NNI (Dasgupta et al. 2000). Hence the following question is natural.

### Problem 1.

What is the complexity of computing the distance between two discrete time-trees?

Although this problem remains open, we make progress towards the solution by establishing several geometric and algorithmic properties of these graphs. First, we demonstrate in the following example that even for caterpillar trees the complexity cannot be derived from that of NNI distance. A *caterpillar tree* (sometimes called a *ladder tree*) has all leaves connected to a single path from the root. A *cherry* is a pair of taxa adjacent to a common internal node in the tree.

### Example 2.

Let *T* be the ultrametric caterpillar tree denoted ((((((1, 2), 3), 4), 5), 6), 7) in Newick notation (Felsenstein et al. 1990) and *R* be the ultrametric tree ((((((1, 4), 5), 6), 2), 3), 7). Then a shortest NNI path is given by first making a cherry (2, 3), then moving the cherry up to the split 1456 | 7, and then resolving the cherry back. In RNNIu this path is not shortest, and one shortest path moves the parents of 2 and 3 up independently.

A set of trees *A* is called conve*x* if for every pair of trees from A, there exists a shortest path between them such that every tree on the path belongs to *A*. Example 2 can be generalized to show that the set of trees of the form (*…* (1, *i*_2_),…,*i_n_*−1), *n*), where {*i*_2_, …,*i*_*n*−1_} = {2,…,*n*−1}, is convex in DtT and is not convex in NNI. Indeed, the example shows that if two leaves have to be moved up, they can be moved independently along a shortest path. The generalization of this statement to an arbitrary number of leaves implies convexity in DtT, while the need to group leaves in NNI as in the previous example shows non-convexity in NNI space. This basic difference in the structure of the two graphs suggests that the complexity of computing the distance is likely to be different.

We proceed by establishing basic geometric properties of discrete time-trees.

### 3.1. Sizes of neighborhoods.

We first bound the number of trees in each graph. As expected, times greatly expand the size of the graphs.

#### Lemma 3

(see (Semple and Steel 2003)). Let |*V*| be the number of vertices in the graph, then |*V*| is equal to

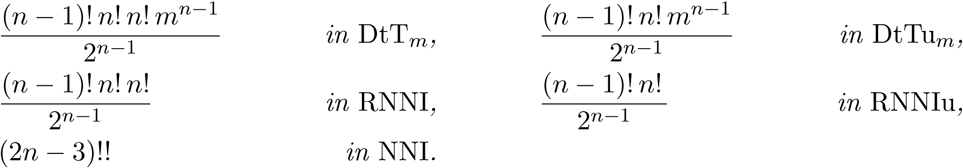

*Proof.* The proof for NNI can be found in (Semple and Steel 2003). The rest follow using similar arguments.

We continue by tightly bounding the sizes of 1-neighborhoods of various discretizations of the space of time-trees. Each has a 1-neighborhood size that is linear in *n* allowing for efficient local traversal and enumeration, a property of importance for phylogenetic algorithms, e.g. tree proposals in MCMC.

#### Lemma 4.

Let *T* beatree on n leaves and deg(*T*)—the number of trees adjacent to T. Then

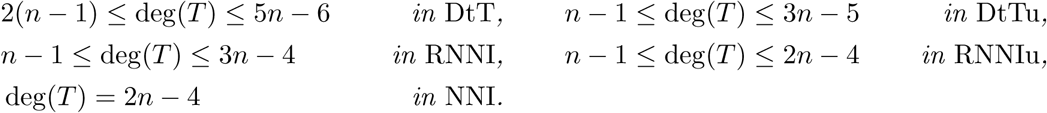

*All bounds are tight.*

*Proof.* We prove the bounds by providing exemplar trees with the specified degrees and explaining why no tree can have a greater (or lesser) degree. Recall that *m* > 1 is the maximal possible length of an event interval in a tree from DtT. The lower bound in DtT is attained by any tree with all event intervals being of length *m*. In this case, deg(*T*) is simply the number of event intervals, since every event interval adds 1 to the total degree of the tree. Other trees have the same or more possible event interval changes, showing that this is a lower bound. The upper bound is attained by a caterpillar tree with all intervals short, every taxon being younger than every divergence event, and both taxa in the cherry being younger than at least one other taxon, that is, a caterpillar tree with internal nodes having ranks *n*, …, 2(*n* − 1) and the ranks of the taxa in the cherry < *n* − 1. In this case, deg(*T*) is bounded by the sum of the 2(*n* − 1) possible interval length changes, at most 2(*n* − 2) NNI neighbors, and at most *n* rank changes that can occur between the *n* taxa and the root of the cherry. In total, deg(*T*) ≤ 2(*n* − 1) + 2(*n* − 2) + *n* = 5*n* − 6. Note that this is an upper bound indeed, as each of the 2(*n* − 2) intervals excluding the most recent can contribute either a rank change or 2 NNI moves. The caterpillar tree described above maximizes the number of intervals that contribute 2 NNI moves and enables the rest of intervals to contribute a rank change.

For ultrametric trees, the number of event intervals is *n* − 1, hence they add *n* − 1 to the degree of the caterpillar tree from interval length changes. The number of intervals on which an NNI move is possible is *n* − 2 for ultrametric trees, hence they contribute 2( *n* − 2) to the degree. In total, this gives the upper bound of 3*n* − 5 for ultrametric trees.

Every NNI move results in two neighbors, so the equality deg(*T*) = 2(*n* − 2) follows for the NNI graph. Indeed, no NNI move can be performed on an edge adjacent to a taxon, exactly two NNI moves can be performed on every edge between internal nodes (the number of which is *n* − 2), and no two trees obtained by an NNI move performed on different edges are identical.

The lower bound in RNNI is attained by the caterpillar-tree where taxa get ranks 0, 1, 3, 5,…, 2*n* − 3 and internal nodes get ranks 2, 4, 6,…, 2*n* − 2. In other words, the divergence events alternate with the taxa in the ranked topology of the tree so that if we parse the tree from the present to the past, we meet the nodes in the following order: taxon, taxon, coalescence, taxon, coalescence, taxon, coalescence, and so on. In this case, intervals bounded by a taxon from below add nothing to the degree of the tree and intervals bounded from below by an internal node add one each, hence *n* − 1 in total. The upper bound for both RNNI and RNNIu is obtained in the same way as for DtT.

For the lower bound in RNNIu, we note that the oldest event interval (the one adjacent to the root) necessarily contributes 2 to the degree. Hence the lower bound can be reached by a tree such that no other interval contributes more than 1. The degree of such a tree is *n* − 1 in RNNIu.

Lemma 4 gives tight bounds for the number of trees in a 1-neighborhood, while the following theorem gives an upper bound for the number of trees in an *r*-neighborhood. The best known upper bound for NNI that we are aware of is 3^*n*−2^2^4*r*^ (Li, Tromp, and Zhang 1996). We obtain a smaller bound for RNNIu and, hence, improve the result for NNI as well.

#### Theorem 5.

The number of trees within distance r from any given tree is at most

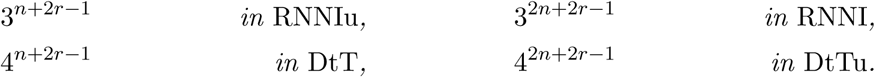

*Proof.* We describe the proof in detail for RNNIu and then explain how to modify the proof for the other three graphs. The proof employs the technique by Sleator, Tarjan, and Thurston (1992) for counting paths in a graph using a *graph grammar*. A graph grammar is a method of encoding each possible graph as a finite set of productions (described more formally below). Given the grammar, the number of possible productions to the power of the encoding length for a given neighborhood radius gives a simple upper bound on the number of trees in a neighborhood. Specifically, Theorem 2.3 from (Sleator, Tarjan, and Thurston 1992), adapted to trees, states the following:

> Let *T* be a tree with *h* nodes, including all internal nodes and taxa, Γ be a graph grammar, *d* be the number of vertices in left sides of Γ, and *k* be the maximum number of nodes in any right side of a production of Γ. Let *R* (*T,*Γ*,r*) be the set of graphs obtainable from *T* by derivations in Γ of length at most *r*. Then |*R* (*T,*Γ*,r*)|≤ (*d* + 1)^*h* + *k*·*r*^.

We first apply Theorem 2.3 from (Sleator, Tarjan, and Thurston 1992) and then improve the bound using specific properties of the RNNIu graph. We note that since our trees are ranked trees, they possess additional structure (ranking) on top of the tree graph structure, hence we will be applying the same style of argument as in (Sleator, Tarjan, and Thurston 1992) but to a somewhat different structure.

We now introduce the *graph grammar* for RNNIu. Since we will apply the graph grammar to modify trees, we use the graph-theoretic terminology here and recall that a tree is a graph. A graph grammar consists of a finite set of productions {*L_i_* →*_i_ R_i_*}, where *L_i_* and *R_i_* are connected undirected edge-end labeled graphs and →*_i_* is a one-to-one map between half-edges of *L_i_* and those of *R_i_*. Here, for every edge in the tree we distinguish between its two ends and refer to them as half-edges—this is necessary to be able to substitute one subgraph by another in a unique way by connecting corresponding half-edges—as in (Sleator, Tarjan, and Thurston 1992). The productions are then applied to the starting tree *T* to derive all possible trees at RNNIu distance up to *r* from *T*. By a *running tree at stage s* we mean the tree obtained after *s* applications of the production rules in the derivation. A production is said to be *ready* at a stage *s* of the derivation if the running tree at stage *s* has a subgraph isomorphic to the left side *L_i_* of the production. A ready production can be applied to the running tree by destroying all nodes corresponding to the left side of the production under the isomorphism and replacing them with the right side *R_i_* of the production. The map →i of the production then says how to reconnect the right side of the production to the half-edges of the running tree that were created after the destruction. The obtained tree is the new running tree for the next stage *s* + 1 of the derivation. See (Sleator, Tarjan, and Thurston 1992) for precise definitions and details.

The graph grammar for RNNIu is the grammar Γ shown on Figure 5. Note that the definition of a graph grammar requires the left sides *L_i_* to be connected and we have a disconnected left side in the third production. We can adapt our graph grammar to fit this requirement by noting that the nodes on the left side of the production must be of consecutive ranks, so we consider those nodes as being adjacent via a second type of adjacency relation by declaring nodes with consecutive ranks to be adjacent. This second type of relation is in addition to the first type of adjacency relation given by the branches of the tree. Furthermore, this second type of adjacency allows us to avoid considering two separate moves for the left and right sister nodes. Indeed, the half-edge labels order the edges to make the distinction between left and right, however since our nodes are ordered via ranking, this ordering of edges is irrelevant. We will further exploit this important property below, where we improve the bound obtained directly from grammar Γ.

**Figure 4.**
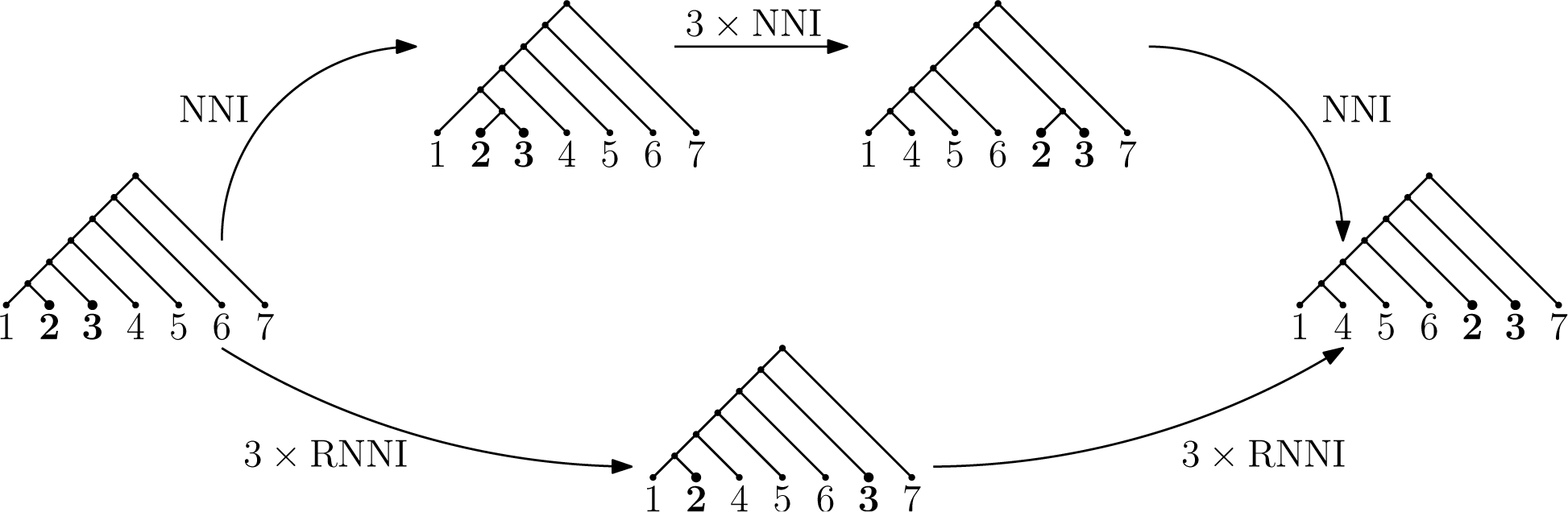
Shortest paths in NNI may not be shortest in RNNI.

**Figure 5.**
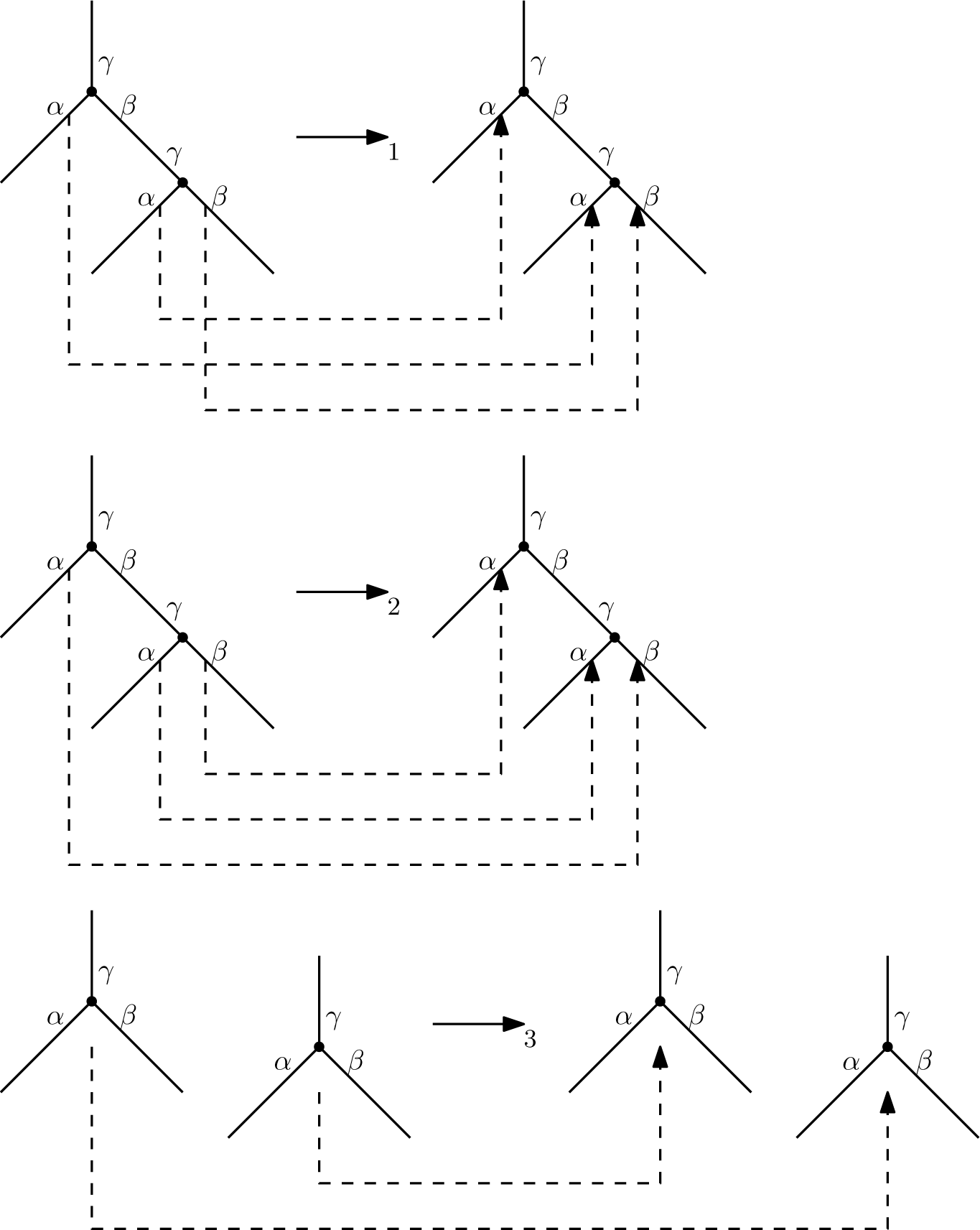
Graph grammar Γ for RNNIu. Maps →_*i*_ are shown by dashed lines. In the first two productions, the edge-ends marked by *β* and *γ* without dashed lines are mapped to each other: top *γ* to top *γ, β* to *β*, bottom *γ* to bottom *γ*. In the last production, edge-ends marked by the same label are mapped to each other. All pairs of nodes on both sides of each production must be of consecutive ranks. The first two productions correspond to an NNI move, while the last production corresponds to rank change.

The number of vertices in left sides of Γ is 6, the maximum number of vertices in any right side of a production of Γ is 2. Since the number of internal nodes of a tree on *n* leaves in RNNIu is *n* − 1, directly applying Theorem 2.3 from (Sleator, Tarjan, and Thurston 1992) we get the bound of 7^*n* + 2*r*−1^.

This bound can be improved following the techniques described in Sections 3.2–3.4 of (Sleator, Tarjan, and Thurston 1992, see also Section 5) by bounding the number of possible productions available at a given time by a number smaller than their total. Every node of a tree in any application of a production of grammar Γ can play either of two roles: top node or bottom node. Hence to indicate that a pair of nodes is ready for being destroyed and replaced using a production, we must specify which node is a top node and which is a bottom node. We use a total of three node-labels (Sleator, Tarjan, and Thurston (1992) call these “labels of vertices”) to identify these two roles and to distinguish which of the two possible NNI moves is applied. We claim that three node-labels *a, b, c* is enough to specify which productions can be applied to a given tree. We must redefine the notion of readiness to verify this claim. We say that a pair of consecutive nodes is *ready* if the node-label of the bottom node is *a* and the node-label of the top node is either *b* or *c*. If two consecutive nodes are ready and not connected by an edge in the tree, then only the rank move is possible. If they are, the type of the move is determined by the node-label of the top node. Hence, this notion of readiness uniquely identifies which of the three productions is applied.

Sleator, Tarjan, and Thurston (1992) use a special node-label, called the “zero label”, to mark nodes that should not be destroyed. They also explain how to eliminate the zero label in various cases. In our case, we can use one of the labels that are already in use, namely node-label *a*. That is, to indicate that a node *v* should never be destroyed, we label the node with an *a*. This labeling will indeed preserve the node *v* because of the following. If *w* is the node directly succeeding *v* at some step of the derivation then *w* can only be marked with an *a* and hence the pair *v, w* is not ready. If *w* is the node directly preceding *v* at some step of the derivation then the pair *w, v* is never ready. Hence *v* is preserved in either case.

Thus, all possible configurations encoded by the six nodes on the left sides of the productions of Γ can be encoded using three node-labels. This implies that the derivation can be encoded by a ternary sequence of length *n* − 1 + 2*r*. Indeed, the first *n* − 1 entries of the sequence are needed to encode all possible node-labelings of the initial tree, plus two entries are needed for each of the *r* applications of productions because every production creates two new nodes, each of which has to be node-labeled. The number of such strings, 3^*n*−1+2*r*^, gives the desired improved bound.

The proof for the other three spaces follows similarly.

For RNNI, we extend the graph grammar Γ by two productions, namely, one for the move when two taxa swap their ranks and one for the move when a taxon and an internal node swap their ranks. Since in both of these productions the only possible type of move is the rank move, two node-labels is enough for these productions. Since the number of (internal and leaf) nodes in an RNNI tree is 2*n* − 1 and the maximum number of nodes on right sides of the productions is still 2, the desired bound is 3^2*n*−1+2*r*^.

For DtTu, we have to extend the grammar Γ by two productions, namely, one for increasing the length of the branch between two nodes in the production and one for decreasing. This increases the number of necessary node-labels by one. Indeed, if a production has a long event interval on the left side, three node-labels are enough to indicate whether the event interval length increases or decreases. Alternatively, if a production has a short event interval on the left side, three node-labels are needed for the top node to indicate the type of move to apply: a length increase move, one of the two possible NNI moves, or a rank swap move. Together, this requires 4 node-labels and the desired bound is 4^*n*−1+2*r*^.

For DtT, we extend the grammar constructed for DtTu by four productions, namely, one for the move when two taxa either swap their ranks or increase the interval length between them, one for the move when two taxa either increase or decrease the interval length between them, one for the move when a taxon and an internal node either swap their ranks or increase the interval length between them, and one for the move when a taxon and an internal node either increase or decrease the interval length between them. All these moves require three node-labels, hence the desired bound is 4^2*n*−1+2*r*^.

Since this theorem bounds the sizes of *r*-neighborhoods by functions of the form *a*^*f*(*n*)+2*r*^, the following corollary bounds the diameters of the graphs under consideration from below.

#### Corollary 6.

Let Δ(*𝒢*), |*𝒢*|, and *δ_r_*(*𝒢*) be the diameter, the number of vertices, and the number of vertices within distance r from any given vertex in graph 𝒢, respectively. If δ_r_(*𝒢*) = *a*^*f*(*n*)+2*r*^ then

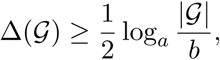

where b = *a*^*f*(*n*)^. In particular, 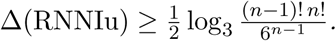

*Proof.* We note that every *r* that satisfies the inequality *δ_r_* (*𝒢*) <|*𝒢*| is smaller than the diameter, that is, Δ(*𝒢*) > *r* for all such *r*. By taking log_*a*_(⋅), the former inequality is equivalent to log*_b_* + 2*r* < log_a_|*𝒢*|, that is, to 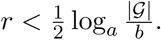 Hence, the desired inequality.

### 3.2. Diameter of the graphs.

Now we estimate the diameter of the graphs from above, complementing the above lower bounds. Recall that Δ(*𝒢*) is the diameter of graph *𝒢*.

#### Theorem 7.

For 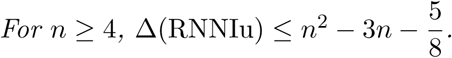

*Proof.* Let *T* and *R* be trees in RNNIu. We show that there exists a path between them of length 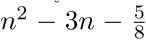 or shorter. We denote the taxa on one side from the root of *T* by *A* and the rest of the taxa by *B*. Let *A* = *A*_1_ ∪ *A*_2_ and *B* = *B*_1_ ∪ *B*_2_ so that the root of *R* splits the taxa into *A*_1_ ∪ *B*_1_ and *A*_2_ ∪ *B*_2_. Note that all *A_i_*’s and *B_i_*’s have to be disjoint. We assume that 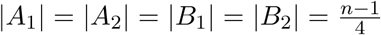 and denote this number by *s*. We will see later in the proof that this assumption does not restrict the generality of our argument.

We construct the path from *T* to *R* proceeding in the following steps, which are illustrated on Figure 6. Let *f*(*n*) be the desired upper bound. For move counts in the remainder of this proof, we will allow the “null move” in which no modification is made.

**Figure 6.**
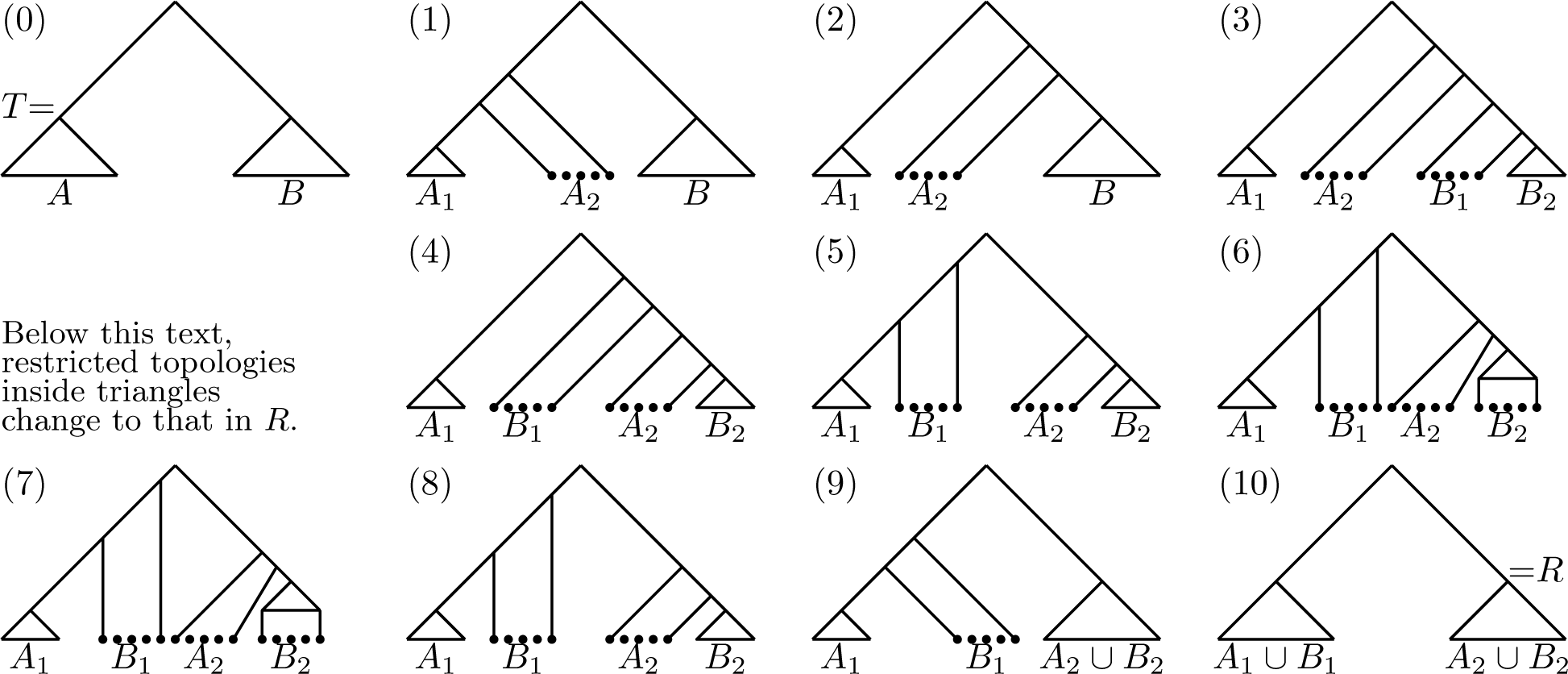
An algorithm to compute a (not necessarily shortest) path between discrete time-trees. Although the trees at steps (6) and (7) look identical, the topologies inside the triangles are different: those at step (6) correspond to the restriction of *T*, at step (7)—of *R*.

1. Move all the nodes adjacent to taxa in *A*_2_ on top of all the other nodes in *T*, apart from the root. The tree restricted to *A*_2_ is then a caterpillar tree. This takes 3*s*^2^ moves.
2. Move the caterpillar tree on *A*_2_ to the other side of the root. This takes 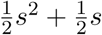 moves.
3. Move all the nodes adjacent to taxa in *B*_1_ on top of all nodes apart from those adjacent to taxa in *A*_2_ and the root. The tree restricted to *B*_1_ is then a caterpillar tree. This takes 2*s*^2^ moves.
4. Swap the caterpillars on *A*_2_ and *B*_1_. This takes *s*^2^ moves.
5. Move the caterpillar tree on *B*_1_ to the other side of the root. This takes 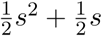 moves.
6. Translate the tree restricted to *B*_2_ up to place it between the trees on *A*_1_ and *A*_2_. This takes *s*^2^ moves.
7. Convert the tree on *A*_1_ ∪ *B*_2_ to the corresponding subtree in *R* in terms of topology and relative ranking of nodes within the subtree. This takes 2*f* (*n/*4) moves.
8. Translate the tree on *B*_2_ down so that the tree on *A*_1_ ∪ *B*_2_ matches the corresponding subtree in *R* in terms of topology and relative ranking of nodes. This takes *s*^2^ moves.
9. Merge the tree on *A*_2_ with the tree on *B*_2_ so that the tree on *A*_1_ ∪ *A*_2_ ∪ *B*_2_ coincides with that in *R*. This takes 2*s*^2^ moves.
10. Merge the tree on *B*_1_ with the tree on *A*_1_ so that the tree coincides with *R*. This takes 3*s*^2^ moves.

In total, we have a recursive equation *f* (*n*) = 14*s*^2^ + *s* + 2*f* (*n*/4).

Assuming that the solution is a polynomial, we see that the polynomial must be of degree 2. It remains to apply the recursive equation to find the coefficients of the polynomial:

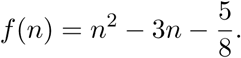

It remains to note that the assumption 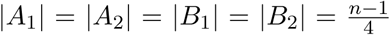 does not increase the distance between *T* and *R*. Indeed, the sets can be chosen so that at most half of the nodes have to cross the root, that is, 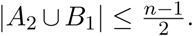. Furthermore, the smaller the size of *A*_2_ ∪ *B*_1_ the shorter the path. Finally, the total length of the path is maximized when |*A*_2_| = |*B*_1_| and |*A*_1_| = |*B*_2_|.

Although we stated and proved the theorem only for the RNNIu graph, the idea can be adapted to the other graphs considered here. However, this adaptation goes beyond the scope of this paper.

We also note that the proof of the theorem provides an efficient algorithm for computing (not necessarily shortest) paths between discrete time-trees. This algorithm can be used for efficient exploration of the space, e.g. via MCMC, where approximate paths can be used to improve mixing by making distant proposals. The algorithm might also be useful to determine if two trees appear to be separated by a valley (e.g. of posterior probability) or if they are rather located on a plateau.

A non-ranked tree has a symmetry associated with every internal node. Those symmetries can be employed to consider only a single representative from equivalent pairs of trees, such as by using tanglegrams (Matsen et al. 2015; Whidden and Matsen 2016), where a *tanglegram* is a graph formed by identifying the leaves of two phylogenetic trees. However, as the following proposition shows, ranked trees are free from all but one of those symmetries, hence the tanglegram approach is not applicable in this case.

#### Proposition 8.

Let *T* be a tree from DtT, DtTu, RNNI, or RNNIu and *σ* a permutation of taxa of *T*. Let *T*_σ_ be the tree obtained from T by permuting the taxa using σ. Then *T* = *T*_σ_ if and only if σ is either the identity permutation or a transposition of a pair of taxa that form a cherry in *T*.

*Proof.* The sufficiency is obvious. For the necessity, assume that *σ* transpose taxa *i, j* such that *i, j* is not a cherry in *T*. This implies that *i* and *j* have different parents. Hence *T* ≠ *T_σ_* because the ranks of the parent of *i* are different in *T* and *T_σ_*.

These observations suggest that the NNI graph is geometrically and algorithmically different from the other four graphs.

### 3.3. Efficient algorithm for generating RNNI graphs

We now introduce an algorithm to compute the RNNI graph on *n* leaves by extending an algorithm from (Whidden and Matsen 2016). The input to our algorithm is a set *S* of RNNI trees in the format described by Gavryushkin and Drummond (2016) and implemented in (Gavryushkin and Drummond 2015) (see also (Semple and Steel 2003)). The two key requirements of the algorithm are the ability to enumerate the neighbors of a given tree in the RNNI graph, and the ability to determine whether a given tree is already a vertex of the graph. Both of these requirements follow from an efficient unique representation for ranked trees.

A node of a tree is given by the set of taxa that are all descendants of the node. A tree is then given by a sequence of nodes ordered by their ranks. Clearly, this obtained representation is unique. For example, the representation of the tree on Figure 1 is ({*A*}, {*C*}, {*D*}, {*B*}, {*E*}, {*A, B*}, {*D, E*}, {*C, D, E*}, {*F*}, { *A, B, F*}, { *A, B, C, D, E, F*}), or ({*A, B*}, {*D, E*}, {*C, D, E*}, { *A, B, F*}, { *A, B, C, D, E, F*}) if we make the tree ultrametric by placing all taxa below the most recent divergence event. Using this representation of trees we construct a map *ν*: *S* → *ω*, where *ω* is the set of non-negative integers. This map allows us to determine whether or not a tree is already a vertex of the graph. This takes linear time if the map is implemented as a trie. We can also enumerate the neighbors of a tree in *O* (*n*^2^)-time using Lemma 4. The algorithm therefore takes *O* (|*S*| * *n*^2^)-time. The high-level steps of the algorithm are as follows. See (Whidden and Matsen 2016) for an analogous proof of correctness.

Construct-RNNI-Graph(*S*)
1. Let *G* be an empty graph.
2. Let *ν* be an empty mapping from trees to integers.
3. Let *i* = 0.
4. For each of the *m* trees:

a. Add a vertex *i* to *G* representing the current tree *T_i_*.
b. Add *T_i_* → *i* to *ν*.
c. For each neighbor *T_j_* of *T_i_*:

(j) If *T_j_* is in dom (*ν*) then add an edge (*i, ν* (*T_j_*)) to *G*.
d. *i* = *i* + 1.

## 4 OPEN PROBLEMS AND CONJECTURES

In the previous section we posed Problem 1, which asks about the complexity of computing the distance in discrete time-tree graphs. This problem is the primary obstacle for actual biological applications of the introduced graphs. The history of research into computational complexity of phylogenetic graphs is very rich and exciting, in particular for the case of the NNI graph, with a number of erroneous results being published over the 25 years it took to settle the complexity. See (Dasgupta et al. 2000) for a detailed discussion of those publications, some of which claimed that NNI distance is NP-hard while others claiming that NNI distance is decidable in polynomial time. The latter claims were mainly based on the so-called Split Theorem which states the following: If a partition of leaves given by an edge is shared between two trees then there exists a shortest NNI path between the trees such that every tree on the path maintains the partition. These claims were finally refuted by Li, Tromp, and Zhang (1996), who proved that the Split Theorem does not hold in the NNI graph. Hence, a natural question that would contribute to understanding Problem 1 is whether or not the Split Theorem holds in the RNNI graph and other time-tree graphs.

### 4.1. Split Theorem.

Li, Tromp, and Zhang (1996) used properties of the diameter of the NNI graph and the sizes of its neighborhoods to provide an example of trees such that every shortest NNI path between the trees fails to maintain a partition of taxa shared by the origin and destination trees. Specifically, they showed that sorting two caterpillar trees simultaneously is more efficient than sorting them independently, provided the size of the trees is large enough. In the following, we conjecture that the ranked versions of NNI graph maintain splits along shortest paths. Our conjecture is based on the fact that RNNI does not satisfy the diameter bounds necessary for the argument in (Li, Tromp, and Zhang 1996) to go through—see Theorems 5 and 7. Indeed, since sorting a caterpillar tree of size *k* in RNNI takes fewer moves than merging two such caterpillar trees and then separating them, the counter-example from (Li, Tromp, and Zhang 1996) does not apply in RNNI. This counter-example using two trees encodes the basic way in which NNI shortest paths are non-trivial: that one can economize by first grouping leaves into bundles, moving the bundles, and then breaking the bundles apart. Because this basic operation does not provide an advantage in the simplest case of two caterpillar trees in RNNI, we do not believe that it will hold for more complex collections of moves.

#### Conjecture 9.

The following is true in all graphs DtT, DtTu, RNNI, and RNNIu, but not in NNI. If a partition of leaves given by an edge is presented in two trees *T* and *R* then the partition is presented in every tree on every shortest path between *T* and *R*.

### 4.2. Computing the distance.

In this section, we bound the number of neighbors of a tree *x* in the discrete time-tree graphs that are closer than *x* to a given tree *y*, **under the assumption that Conjecture 9 holds**. We denote trees by lowercase letters to stress the fact that we are considering graphs as metric spaces in this section. We show that the maximum fraction of neighbors that tend closer to *y* grows linearly with respect to *d* (*x, y*). When the graph is seen as a metric space, this property is important for the study of the curvature of the space as well as convergence properties of random walks over the space. This will be useful in future studies analogous to (Whidden and Matsen 2016). Furthermore, the result provides a further insight into possible approaches to Problem 1.

#### Theorem 10.

Assume that Conjecture 9 holds. Let *x* and *y* be two trees and *N* (*x*)—a one-neighborhood of *x*. Then the number of trees *u* ∊ *N* (*x*) such that *d* (*u, y*) ≤ *d* (*x, y*) is at most

1. 3*d* (*x, y*) *in* RNNI,
2. 4*d* (*x, y*) *in* DtT_2_,
3. 5*d* (*x, y*) *in* DtT_*m*_ *for m >* 2.

*Proof.* We first prove the statement for DtT_2_. Let *U* be the set of neighbors *u* of *x* such that *d* (*u, y*) ≤ *d* (*x, y*), as stated in the theorem. We partition *U* into three sets of trees—*I*, *R*, and *L*— obtained from *x* by a single NNI move, rank change, or length change, respectively. We then prove the theorem by bounding the size of each partition by 2*d* (*x, y*), *d* (*x, y*), and *d* (*x, y*), respectively.

We first consider some basic properties of minimal length paths between *x* and *y*. Observe that at most *d* (*x, y*) bipartitions differ between *x* and *y*, as each NNI operation replaces one bipartition with another (and rank and length changes do not modify bipartitions). Now, Conjecture 9 implies that no minimal length path from *x* to *y* will replace a bipartition that is common to *x* and *y*.

We are now ready to bound the sizes of each partition *I, R*, and *L*. First, consider the subset *I* of closer neighbors obtained via NNI moves. By our observations above, |*I*| is bounded by the number of NNI operations that modify one of the at most *d* (*x, y*) bipartitions of *x* that are not a bipartition of *y*. There are two neighbors of *x* that lack any given bipartition (obtained by moving either the left or right subtree located below the bipartition). Therefore, |*I*| ≤ 2*d* (*x, y*).

Second, consider the subset *R* of closer neighbors obtained via rank change moves. These operate either on one of *x*’s unique bipartitions or a bipartition common to *x* and *y* that differs in rank. As observed above, the number of unique bipartitions is bounded by the maximum number of NNI moves on any minimal *x* to *y* path. Any such path must fix the ranks of each common edge. In other words, if *r*_1_ is the number of trees in *U* which are obtained from *x* by a rank move corresponding to a common edge and *r*_2_ is the number of trees in *U* which are obtained from *x* by a rank move corresponding to a unique edge of *x*, then *r*_1_ + *r*_2_ ≤ *d* (*x, y*), because the *r*_1_ moves have to be done along every shortest path from *x* to *y*. Thus, the total size of *R* is bounded by *d* (*x, y*).

The bound of *d* (*x, y*) on the number of length changes that can be applied to move *x* closer to *y* follows similarly. Therefore, there are at most |*I*| + |*R*| + |*L*|≤ 2*d* (*x, y*)+ *d* (*x, y*)+ *d* (*x, y*) = 4*d* (*x, y*) trees *u* ∊ *N* (*x*) such that *d* (*u, y*) ≤ *d* (*x, y*).

The statement for the other two graphs follows similarly: for RNNI we need to count only for trees from *I* and *R*, for DtT we will have to add 2*d* (*x, y*) instead of *d* (*x, y*) for *L*.

We conclude by noting that most of the results in this paper can be generalized to the class of sampled ancestor trees (Gavryushkina, Welch, et al. 2014), where DtT trees are enriched by an extra event called *sampled ancestor*—an internal node marked by a taxon with exactly one child. However, currently only a few methods of sampling the space of sampled ancestor trees are known (Gavryushkina, Heath, et al. 2016) and the geometry of the space is poorly understood (Gavryushkin and Drummond 2016), hence we do not include the generalization here.

## 5. DISCUSSION

In this paper, we have introduced a hierarchy of discrete time-trees that naturally approximates the full space of phylogenetic time-trees. The first two levels of the hierarchy are the well-known classes of rooted phylogenetic trees and ranked phylogenetic trees. We extend this previous work by considering a novel graph structure, the RNNI graph, on the set of ranked trees. In the same way that the NNI graph is the discrete component of the full space of phylogenetic trees, the RNNI graph is the simplest discrete component of the full space of time-trees. Hence, the introduced hierarchy fits and refines naturally the classical picture of phylogenetic graphs.

Surprisingly, the geometry of RNNI and NNI graphs differ in many aspects including the diameter, the sizes of neighborhoods, and the convexity of caterpillar trees. This suggests that the computational complexity of computing distances in the two graphs is likely to be different, at least at the level of certain subgraphs such as the restriction to caterpillar trees. Although the size of discrete ranked graphs are much larger than their NNI graph counterparts, ranks provide an inherent structure that may serve as a useful avenue of attack for future algorithms.

We introduce and make some initial progress on the computational complexity of calculating the distance between two time-trees (Problem 1). This is an important step for further progress in theoretical research as well as computational and biological applications of time-trees. In particular, a better understanding of this problem would have the following applications.

### 5.1. Efficient tree search algorithms

Some of the most popular phylogenetic time-tree inference methods utilize the Markov Chain Monte Carlo (MCMC) algorithm to sample a probability distribution over the space of time-trees. The most efficient proposals on time-trees will take into account both local and global geometry of the corresponding space. Our hierarchy of discrete time-trees offers a natural discrete structure of the space used in such tree proposals.

Thus far, the most widely used modifications of trees are NNI, SPR, and their variants. The graph distance inherited from these modifications is known to be NP-hard to compute, and although sometimes NP-hard problems can be solved efficiently for practically interesting data sets, this is not the case for these phylogenetic graphs of ranked trees, where the size of tractable problems is an order of magnitude smaller than those that routinely arise in practice. Hence, a computationally tractable phylogenetic graph is a highly relevant and desirable object for the field, and we believe that the RNNI graph should have this property.

### 5.2. Convergence of phylogenetic algorithms

A tractable phylogenetic distance would greatly assist methods to assess convergence of Bayesian tree inference algorithms based on MCMC and related methods. In other areas of statistics, distance methods are widely used to test for convergence, but the computational complexity of phylogenetic distances makes those methods applicable to only very moderate data sizes. Phylogenetic distances are especially suitable for such applications because they correspond to the distance inherited from the inference algorithm, being based on the same sorts of tree rearrangements.

Another important property for phylogenetic applications is to compute the number of trees that are “equally good” under certain criterion. This intuition is formalized by Sanderson, McMahon, and Steel (2011) in the notion of a terrace. To compute the terrace, an efficient tree metric is of crucial importance.

